# EZ Clear for simple, rapid, and robust mouse whole organ clearing

**DOI:** 10.1101/2022.01.12.476113

**Authors:** Chih-Wei Hsu, Juan Cerda, Jason M. Kirk, Williamson D. Turner, Tara L. Rasmussen, Carlos P. Flores Suarez, Mary E. Dickinson, Joshua D. Wythe

## Abstract

Tissue clearing for whole organ cell profiling has revolutionized biology and imaging for exploration of organs in three-dimensional space without compromising tissue architecture. But complicated, laborious procedures, or expensive equipment, as well as the use of hazardous, organic solvents prevents the widespread adoption of these methods. Here we report a simple and rapid tissue clearing method, EZ Clear, that can clear whole adult mouse organs in 48 hours in just three simple steps. Samples stay at room temperature and remain hydrated throughout the clearing process, preserving endogenous and synthetic fluorescence, without altering sample size. After wholemount clearing and imaging, EZ Cleared samples can be further processed for downstream embedding and cryosectioning followed by standard histology or immunostaining, without loss of endogenous or synthetic fluorescence signal. Overall, the simplicity, speed, and flexibility of EZ Clear make it easy to adopt and apply to diverse approaches in biomedical research.

## INTRODUCTION

Over the past 30 years, the development of confocal microscopes that can image large samples at cellular resolution, combined with powerful increases in computing and the ability to handle large volumes of data, have unleashed an explosion in three-dimensional (3D) visualization of organ structures at both a macro and cellular scale ^1,2^. A key to this revolution has been a simultaneous deluge of tissue clearing protocols driven by advances in optical physics and chemical engineering ^1–3^. The development of both organic solvent-based clearing methods like BABB ^4^, 3DISCO ^5,6^, iDISCO ^7^, Ethanol-ECi ^8^, PEGASOS ^9^, Fast 3D ^10^, and aqueous-based techniques such as Sca*l*e ^11,12^, CLARITY ^13^, PACT-PARS ^14^, CUBIC ^15^, and Ce3D ^16^ have been successfully leveraged in various biological model systems. As light microscopy-based deep imaging of larger intact organs or samples are usually limited by the scattering of light, advances in tissue clearing allows researchers to examine tissues in their native 3D state, by imaging modalities with different volume capacities and resolution, ranging from optical projection tomography (OPT), confocal, multiphoton, to cutting edge lightsheet fluorescence microscopy (LSFM)-based imaging ^17–22^.

Although each of the aforementioned strategies has their own unique merits ^1–3^, several hurdles remain for the field in terms of developing and adopting a rapid, simple, and robust tissue clearing method. While aqueous-based methods may preserve fluorescence from endogenous transgenic reporters, and can be easily imaged on majority of the existing imaging platforms, they often require either extended incubation periods (days to weeks) or complicated, laborious procedures, as well as special clearing equipment ^1,3,23^. For example, hydrogel-scaffolding based CLARITY clearing provides a robust and controllable workflow to clear tissue in the aqueous environment, but the complicated steps and considerable technical investment represent substantial hurdles that prevent many researchers from adopting this methodology ^13^. Moreover, as the hydrogel scaffolding is based on covalently conjugating proteins in a polyacrylamide matrix, precise control of tissue fixation with paraformaldehyde and thermal crosslinking of polyacrylamide is required. Otherwise, the overall strength and pore size of the hydrogel-tissue matrix varies experiment to experiment during SDS-mediated electrophoretic removal of lipids, limiting reproducible, robust results ^3,13,24^. In addition, variations in clearing efficiency for different organ systems based on tissue composition and volume also pose a significant challenge for researchers, as extensive optimization is required to tailor this method to individual research projects.

In contrast, organic solvent-based clearing methods are simple, fast, efficient, and do not require specialized homemade or commercial equipment. However, challenges in sample handling and imaging tissues in the hazardous, environmentally toxic solvents, combined with the rapid decline of endogenous fluorescent signals, limit its applicability ^1,3,21,23,24^. Organic solvent-based clearing strategies typically use corrosive and combustible organic solvents to match the refractive index (RI) of dehydrated tissues to reduce light scattering and render tissue optically transparent, which limits downstream imaging to microscope systems with solvent-resistant chambers and objectives. Critically, while samples cleared by aqueous-based strategies can be subsequently processed for conventional histology and immunostaining ^25^, the ability to process samples after solvent-based clearing and proceed with cryosection to further investigate tissue sections with histology or immunofluorescence staining are yet to be seriously explored, limiting the uses of precious tissue samples in these approaches.

Here, we present a simple and rapid tissue clearing method, EZ Clear, that can effectively clear whole adult mouse organs in 48 hours in three simple steps: EZ Away, EZ Wash, and EZ View. EZ Clear combines the advantages of solvent-mediated lipid removal with highly water-miscible tetrahydrofuran (THF) and renders sample transparent in an aqueous, high RI (n = 1.518) sample mounting and imaging solution compatible with most microscopy platforms. Because samples are submerged in an aqueous environment during the entire protocol, no significant changes in size occur within the tissue. Herein we use EZ Clear to render multiple adult mouse organs optically clear (brain, eye, heart, lung, liver, pancreas, kidney, testis) in 2 days. Our results demonstrate that EZ Clear processed samples not only retain endogenous fluorescence from transgenic reporters to allow 3D whole organ lightsheet imaging, but EZ Cleared samples can also be processed after wholemount imaging for cryosectioning and histology or immunofluorescence staining for further analysis. In summary, EZ Clear is a simple, robust, and easy to adopt whole organ clearing technique that can be applied to various sample volumes and utilized across most common imaging platforms.

## RESULTS

### EZ Clear: simple, rapid, and effective

EZ Clear effectively clears adult mouse organs in three simple steps within 48 hours at room temperature (Figure 1A). Step 1 is to immerse the fixed sample in EZ Away solution, which consists of 50% (v/v) tetrahydrofuran (THF) in sterile Milli-Q water. THF is highly water-miscible, easily infiltrates biological samples, solubilizes lipids, and minimizes fluorescent quenching ^5,26,27^. As EZ Away is 50% water, delipidation of tissue occurs while the organ remains in an aqueous environment. Following lipid removal, the sample is incubated in EZ Wash (sterile Milli-Q water) for four hours to remove any remaining THF from the tissue. The tissue is then rendered transparent by immersing it in aqueous EZ View sample mounting and imaging solution at room temperature for 24 hours. EZ View solution is 80% (v/v) Nycodenz, 7M urea, and 0.05% sodium azide prepared in 0.02 M phosphate buffer (pH 7.4) and has an RI of 1.518. Compared to brains fixed in only 4% paraformaldehyde (PFA), EZ Clear not only renders the whole brain as transparent as 3DISCO and FAST 3D, but EZ Clear also has the simplest procedure (3 steps) and shortest processing time (48 hours) (Figure 1B, S1A). In addition, because the sample is maintained in an aqueous environment throughout the entire EZ Clear process, tissue size remains constant throughout the protocol, with no significant change after clearing (Figure 1B, n=4 for each condition, one-way ANOVA, size change ratio = 1.072 ± 0.062). Conversely, other approaches either significantly shrank (Fast 3D: 0.776 ± 0.025 and 3DISCO: 0.593 ± 0.013) or expanded (X-CLARITY: 1.608 ± 0.049) the brain (Figure 1B and Figure S1B).

**Figure 1.**
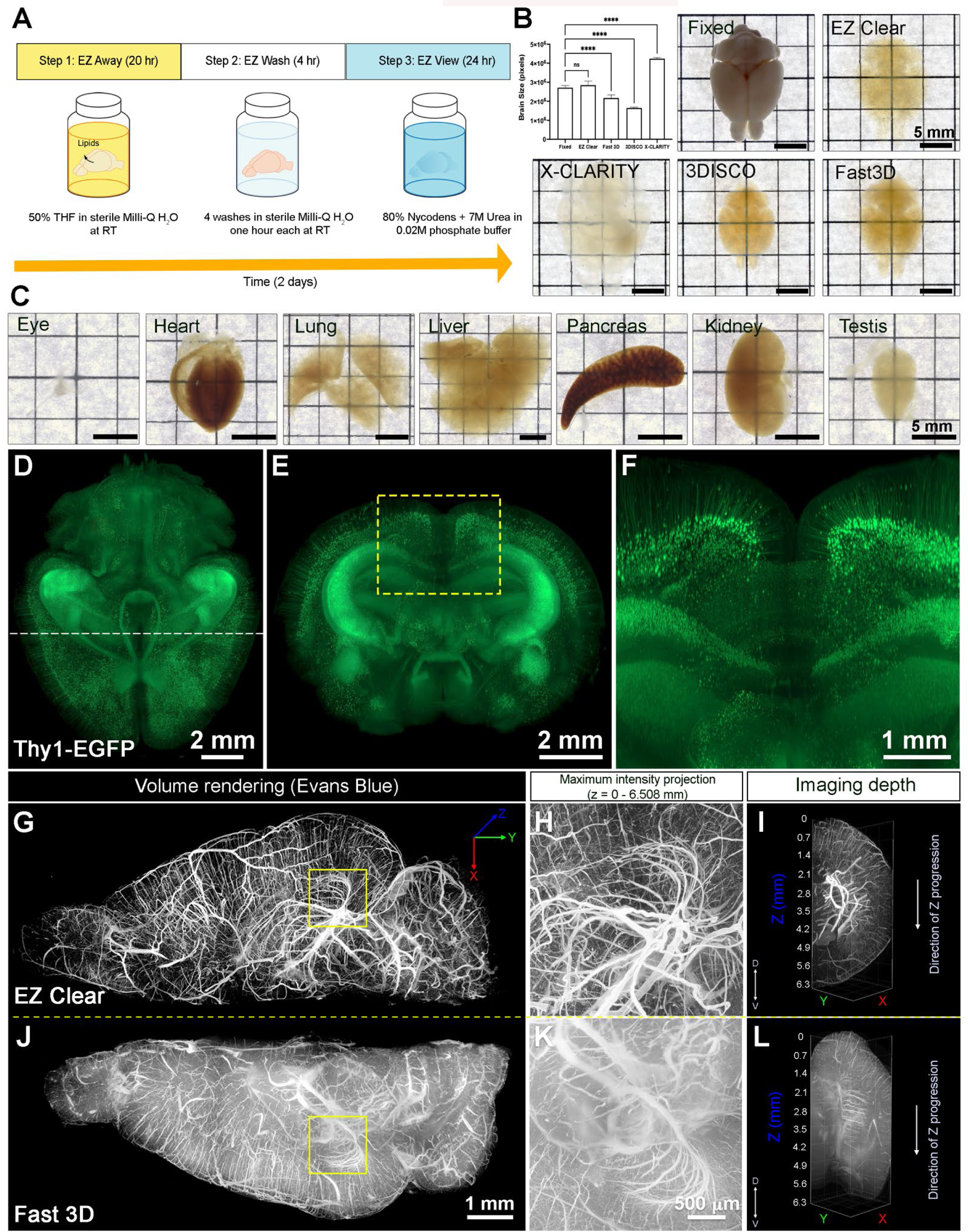
EZ Clear is a simple, rapid, and efficient tissue clearing process. (A) Procedure for EZ Clear tissue processing. (B) Comparison of volume changes of adult mouse brains (9 weeks old) and bright field representative images of fixed in 4% PFA and not cleared, or fixed and cleared using EZ Clear, Fast 3D, 3DISCO, and X-CLARITY cleared brains. (C) Adult mouse organs (eye, heart, lung, liver, pancreas, kidney, and testis) cleared by EZ Clear protocol. (D-F) Volume rendering of wholemount, 4-month-old *Thy1-EGFP-M* mouse brain cleared with EZ Clear and imaged by LSFM across the total imaging depth of 5 mm, dorsal to ventral at (D) dorsal view and (E and F) sectioned digitally at the transverse (coronal) axis. (G – L) Lightsheet imaging of the right or left hemisphere of a mouse brain perfused with Evans blue dye and then cleared by EZ Clear (right hemisphere, G-I) and Fast 3D (left hemisphere, J-L) shows that EZ Cleared tissue has a greater imaging depth with cleaner signal and less light scattering.

EZ Clear is also applicable to other mouse organ systems, not just the mouse brain, and achieves high tissue transparency (Figure 1C). In addition, signal from endogenous fluorescent transgenic reporter lines is preserved by EZ Clear, as a brain from a *Thy1-EGFP* transgenic neuronal reporter line that was processed with EZ Clear retained robust signal when imaged by lightsheet microscopy across the total imaging depth of 5 mm, dorsal to ventral (Figure 1D-F and Movie S1). To further assess the clearing efficiency and sample transparency of EZ Clear, we compared samples cleared by two THF-based clearing methods, EZ Clear and Fast 3D, following administration of a synthetic, exogenous fluorescent dye that labels the endothelium. When Evans blue, a non-cell permeable dye, binds to albumin, it undergoes a conformational shift that produces fluorescence in the far-red spectrum (excitation at 620 nm, emission at 680 nm) ^28,29^. A cost affordable alternative to expensive dyes, such as fluorescent tomato lectin, Evans blue can thus be used to robustly label the endothelium following intravenous administration ^30^. Adult mice perfused with Evans blue were euthanized and the brains were harvested and fixed in 4% PFA overnight at 4°C. The brains were then bisected at the midline along the anterior-posterior axis and the left hemisphere was cleared with Fast 3D, while the right hemisphere was cleared with EZ Clear. Both hemispheres were imaged independently on a Zeiss Lightsheet Z1 from the dorsal to ventral side with an EC Plan-Neofluar 5x/0.16 air detection objective at a resolution of 1.829 μm laterally and 3.675 μm axially (Figure 1G-L and Movie S2). The right hemisphere, processed with EZ clear, was equilibrated and imaged in EZ View solution (RI = 1.518), while left hemisphere, cleared with Fast 3D, was equilibrated and imaged in Fast 3D imaging solution (RI = 1.512). Although both EZ Clear and Fast 3D can render the tissue highly transparent, lightsheet imaging revealed that EZ Clear enables imaging depths up to 6.5 mm from dorsal to ventral sides, with clean and sharp signals (Figure 1G-I), whereas light scattering increased when imaging deeper into the hemisphere processed using Fast 3D (Figure 1J-L). Thus, in 48 hours and three simple steps, EZ Clear achieves transparencies comparable to 3DISCO, but without significantly changing sample size, while preserving endogenous and synthetic fluorescence, and it is compatible with aqueous-based imaging platforms. Additionally, EZ Clear is compatible with extensive imaging depths and induces minimum light scattering.

### Tissue RI matching with aqueous EZ View sample mounting and imaging solution

To maintain cleared samples in an aqueous environment following delipidation by EZ Away, we compared sample mounting and imaging buffers for rendering tissue optically transparent. While the refractive index of water is estimated to be 1.33, the RI of the soft tissue is between 1.44 and 1.56, and the RI of dry tissue is approximately 1.50 ^21,31,32^. In solvent-based clearing methods (e.g., 3DISCO and BABB), dehydrated and delipidated samples are equilibrated in high RI solvents, such as DBE (RI = 1.56) and BABB (RI = 1.55) to reduce light scattering. In aqueous-based clearing methods, solutions like RIMS (80% (v/v) Nycodenz, RI = 1.46) and sRIMS (80% D-sorbitol, RI = 1.43) are suitable for mounting and imaging of SDS-mediated delipidated tissue, although they are lower than the ideal RI of 1.52-1.56 ^14,33^. Given the aqueous nature of EZ Clear, we first tested whether RIMS or sRIMS were compatible with EZ Away. Somewhat surprisingly, EZ Away treated samples immersed in either RIMS (Figure 2C) or sRIMS (data not shown) were not rendered transparent. Thus, we pursued the novel aqueous solutions with an RI over 1.50. Previous studies have shown that urea can hyperhydrate samples and enhance tissue transparency ^11,15,34^. To test the effect of combining urea and Nycodenze on RI and clearing efficiency under aqueous conditions, we gradually increased the urea concentration in RIMS and measured changes in RI (Figure 2A). The solution was saturated with 8M of urea in 80% Nycodenz, but the RI reached the highest value (RI = 1.518 α 0.0003) with 7M urea in 80% Nycodenz at room temperature. We then assessed sample transparency by comparing an EZ Away treated brain immersed and equilibrated in either PBS (RI = 1.332), RIMS (80% Nycodenz, RI = 1.463 α 0.0019) or EZ View (80% Nycodenz and 7M urea, RI = 1.518) (Figure 2B-D). While the tissue remained opaque in PBS (Figure 2B), the transparency of the EZ Away treated brain only showed minor improvement following immersion and equilibration in RIMS for 24 hours (Figure 2C). However, the EZ Away treated brain immersed and equilibrated in EZ View solution for 24 hours was comparable to samples cleared using 3DISCO (compared Figure 2D and Figure 1B). We next examined the effect on imaging depth of EZ Away processed samples cleared in different RI matching solutions. Adult mice perfused with 649 *Lycopersicon esculentum* (tomato) lectin fluorescently conjugated with DyLight (lectin-649), which labels the endothelium, were euthanized and the brains were harvested and immersion fixed in 4% PFA overnight at 4 °C. After fixation, lectin-649 labeled brains were treated with EZ Away and EZ Wash and then either equilibrated and imaged in RIMS or EZ View by lightsheet fluorescent microscopy (Figure 2E-J). Not only were brains equilibrated in EZ View were more transparent than those equilibrated in RIMS (Figure 2C and D), but they also featured imaging depths over 6 mm through the dorsal-ventral axis of the brain (Figure 2F and J). Color-coded depth projection stepping at 1 mm intervals showed that while the signal from lectin-649 starts to scatter 1 to 2 mm deep into RIMS treated tissue (Figure 2G), the signal remains sharp and clean for the brain equilibrated and imaged in EZ View up to a depth of 6 mm (Figure 2J), with all vessels easily distinguishable from the dorsal to ventral side (Figure 2K-M). Critically, the EZ Clear samples remain in aqueous environment throughout the entire clearing and imaging process, and no corrosive and combustible organic solvents or immersion oil are needed for RI matching or imaging.

**Figure 2.**
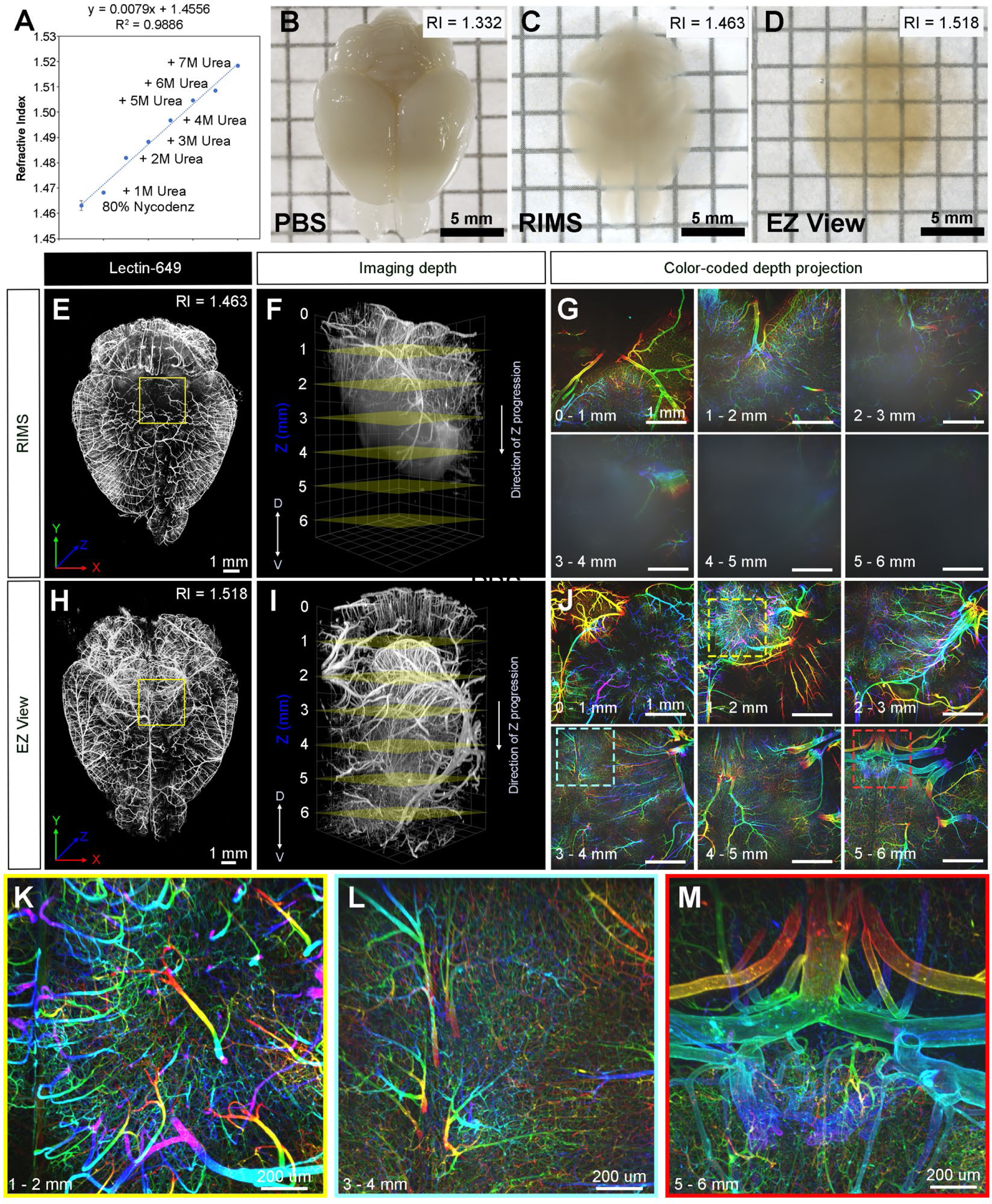
Aqueous EZ View mounting and imaging buffer with high/adjustable refractive index. (A) Refractive index (RI) of 80% Nycodenz increases linearly with increasing concentrations of urea. (B-D) Comparison of transparency of adult mouse brains processed with EZ Away and EZ Wash and then equilibrated in (B) PBS (RI = 1.332), (C) RIMS (RI = 1.463), and (D) EZ View (RI = 1.518) for 24 hours at room temperature. (E-J) Comparison of 3.5 months old mouse brains perfused with far-red fluorescent lectin, then treated with EZ Away/EZ Wash and equilibrated in (E-G) RIMS and (H-J) EZ View and imaged by LSFM. (E and H) Comparison of volume rendered whole brains, transverse view, and (F and I) across the imaging axis starting from dorsal to ventral, and (G and J) color coded depth projection at 1 mm intervals beginning from dorsal (0 mm) to ventral (6 mm) side. (K-M) Representative color-coded depth projection images of EZ View equilibrated and lightsheet imaged sample at (K) 1 to 2 mm, (L) 3 to 4 mm, and (M) 5 – 6 mm.

### EZ Cleared and imaged samples can be further processed for cryosectioning, histology, and immunofluorescence staining

To further determine the downstream utility of EZ Cleared samples, we next explored the possibility of processing EZ Cleared and wholemount imaged samples for embedding and cryosectioning. For these studies, we examined brain tumor formation in a native murine model of glioma, as before ^35^. To induce glioma, the lateral ventricle of E16.5 mouse embryos were injected with a DNA cocktail consisting of 3 plasmids: (1) a single pX330-variant construct encoding 3xFlag-NLS-Cas9-NLS, along with three U6 promoter cassettes upstream of guide RNAs targeting the tumor suppressor genes *Nf1, Trp53*, and *Pten*, (2) a *piggyBac* (PB) helper plasmid with the glial- and astrocyte-specific promoter, *GLAST (EAAT1)*, driving expression of PB transposase, and (3) a PB cargo fluorescent EGFP reporter vector to indelibly label all tumor cells and their descendants ^35^. Following injection of the DNA cocktail, the embryos were electroporated to allow uptake of the constructs, then the embryos were placed back in the maternal cavity. These in utero electroporated (IUE) animals with tumor suppressor deficient cells were then birthed normally and were collected at postnatal day 104 (P104) for perfusion with lectin-649, and then the brains were harvested, fixed, and processed with EZ Clear and wholemount imaged with lightsheet to reveal the distribution of GFP^+^ tumor cells and the lectin-649 labeled vasculature (Figure 3A and B). At a macro level via whole brain imaging, the GFP^+^ tumor cells can be identified not only as clusters (Figure 3A and B), but also as single cells (Figure 3C and D, Movie S3). After imaging, the glioma containing samples were immersed in 1X PBS overnight to remove the EZ View solution, then processed through a graded series of solutions with increasing sucrose concentrations and then embedded in OCT for cryosectioning. EZ cleared P104 glioma brains were then cryosectioned for direct immunofluorescence and also for Hematoxylin and Eosin (H&E) staining to visualize tissue architecture. After cryosectioning, tissue sections were mounted and inspected on both fluorescence and confocal microscopes to evaluate whether the fluorescence from the transgenic EGFP reporter and synthetic fluorescent lectin dye were preserved. Sections mounted in EZ View solution became highly transparent compared to those remained in PBS or mounted in Prolong Glass Antifade medium (Figure 3E-G). Sections mounted in EZ View were compatible with both fluorescence (Figure 3H) and confocal imaging (Figure 3I-J), as signals from GFP^+^ tumor cells and perfused Lectin-649 vessels were well preserved. EZ Cleared, imaged, and cryosectioned tissues were also compatible with conventional histology, as evidenced by robust H&E labelling of tissues (Figure 3K-M) comparable to tissues fixed using only 4% PFA (Figure S2).

**Figure 3.**
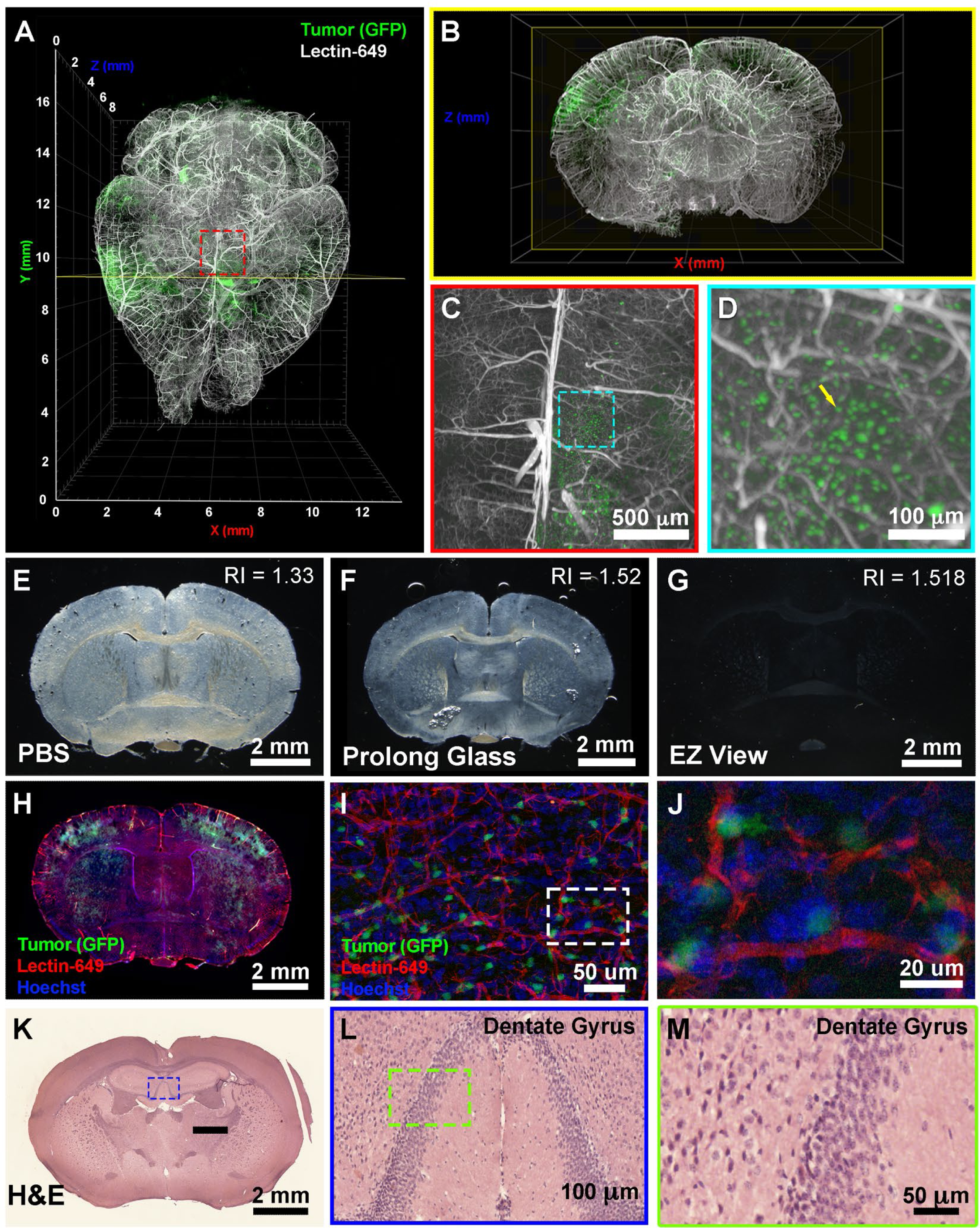
EZ Cleared and imaged samples can be further processed for downstream cryosectioning, histology, and immunofluorescence staining. (A) Volume rendering of wholemount lightsheet imaged, EZ Cleared P104 glioblastoma multiforme (GBM) mouse brain with GFP^+^ tumor cells and lectin-649 labeled vasculature at (A) dorsal view and (B) sectioned digitally at the transverse (coronal) axis. GFP^+^ tumor cells can be identified in large clusters (A and B), as well as sparse single cells (C and D) from wholemount imaged data. (E-G) EZ Cleared and imaged brains processed for cryosectioning in the coronal plane, and then mounted in (E) PBS, (F) Prolong Glass Antifade medium (n = 1.52), or (G) EZ View. Sections mounted in EZ View is compatible with fluorescence (H) and confocal (I and J) imaging and the signals from GFP^+^ tumor cells and fluorescent lectin labeled vasculature are preserved. Tissues processed for EZ Clear are also compatible with downstream histological applications, as cryosections stained with Hematoxylin and Eosin (H&E) yielded robust labelling of nuclei and cytoplasm (K-M).

Next, we explored whether EZ Cleared and cryosectioned tissues are compatible with indirect immunofluorescence. Free-floating sections from brains harboring EGFP^+^ glioma cells that were perfused with far-red fluorescent lectin were processed through a simple, six step immunofluorescence staining protocol (Figure 4A). Sections were stained with CD31 (to label endothelial cells), GFAP (astrocytes), smooth muscle α-actin conjugated with Cy3 (αSMA-Cy3) (smooth muscle enwrapped blood vessels), and class III β-tubulin (e.g., Tuj1, to label neurons), and mounted with EZ View solution for confocal imaging using an 880 Airyscan confocal microscope. Confocal imaging reveals that in addition to preserving the fluorescence of GFP+ tumor cells and perfused Lectin-649 labelled endothelial cells, each of these distinct antigens were robustly detected by indirect immunofluorescence following EZ Clearing and cryosectioning (Figure 4B – D, Movie S4). These results demonstrate that EZ Clear processed and imaged tissue not only can be interrogated at a macro level by whole organ 3D imaging using LSFM, but processed tissue can also be further used for cryosectioning and investigated at the cellular level via either histological or immunofluorescent staining.

**Figure 4.**
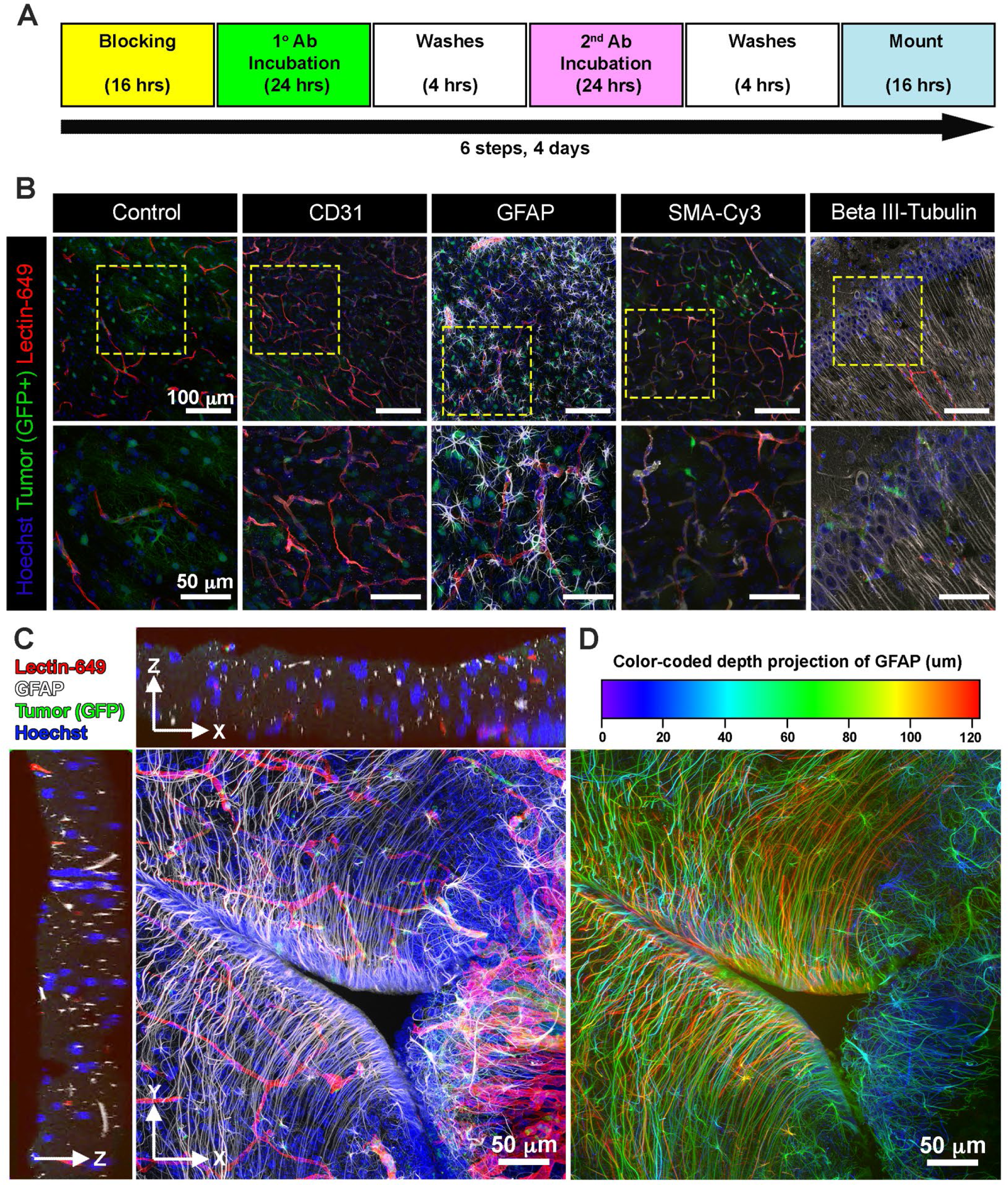
Cryosectioned EZ Cleared and imaged samples is compatible with immunofluorescence staining. (A) 6 step immunostaining procedure for processing tissues after EZ Clearing and imaging. (B) Coronal plane cryosections from EZ Cleared and imaged P104 glioblastoma multiforme (GBM) mouse brains with GFP^+^ tumor cells and lectin-649 labelled vessels were unstained (Control), or immunostained to detect CD31 (endothelial), GFAP (astrocyte), smooth muscle α-actin (smooth muscle cells), and β-III tubulin (neurons). Stained sections were mounted on slides with EZ View and imaged on an 880 Airyscan FAST confocal microscope at 20X. (C) Orthogonal and (D) color-coded depth projection views of GFP^+^ tumor cells and lectin-649 labelled vessels section labeled with GFAP and Hoechst. The immunostained GFAP signal was constantly throughout the 100 μm section.

## DISCUSSION

In this work, we developed a simple, rapid, and robust tissue clearing procedure, EZ Clear, that renders entire adult mouse organs transparent within 48 hours, without the need for special equipment or toxic organic solvents. By combining a water-miscible solvent that rapidly infiltrates and dissolves lipids within a tissue, and an aqueous high refractive index sample mounting medium that minimizes light scattering and renders tissues optically transparent, EZ Clear makes whole organ clearing and high-resolution imaging simple and compatible with a variety of imaging needs.

EZ Clear has several advantages over other clearing methods. First, unlike electrophoretic-based CLARITY clearing, EZ Clear does not require a costly up front financial investment in specialized equipment, as its simplicity and robustness make it easy to adopt and apply immediately in any standard molecular biological laboratory. Additionally, samples require only little attention during processing, and this simple, three step protocol requires minimal time and effort investment, as it yields cleared tissue within 48 hours. Furthermore, by using an aqueous sample mounting and imaging solution with a high RI (1.518), delipidated samples remain hydrated while being imaged, preserving endogenous fluorescent signals, and making it compatible with most fluorescent imaging platforms. Samples can also be imaged in EZ View without the requirement of immersion oil, further simplifying the procedure of processing cleared and imaged sample for downstream process. We also note that EZ View solution can be used as a sample mounting and imaging medium for cryosectioned tissue. Finally, the ability to process EZ Cleared and imaged samples for downstream cryosectioning, histology and immunofluorescence staining creates a powerful workflow to further interrogate tissues. While whole organ immunostaining is possible with currently established protocols, such as iDISCO and CUBIC-HV ^7,36^, we propose that whole organ imaging with EZ Clear, followed by interrogation of thicker sections via cryosectioning and immunohistochemistry-based indirect immunofluorescence, allows researchers to maximize the yield of precious tissue samples, while minimizing the costs associated with excessive antibody use and saves time by allowing one to focus on a specific region of interest.

A shortcoming of the present study is that we have not yet tested the method on larger samples, or tissues from other species such as rat, pig, or humans, nor did we examine mouse tissues at any younger ages (e.g., embryogenesis). Future testing on the volume of EZ Away required, and incubation time necessary, scaled to various sample sizes with varying lipid contents will be necessary for further optimization of the technique, but we anticipate that this method will readily work with samples smaller than the adult mouse organs shown in the present study.

In summary, EZ Clear renders adult mouse organs optically transparent in 48 hours in just three simple steps using standard, off the shelf reagents in an aqueous-based tissue clearing methodology. EZ Clear is a simple, robust, and easy to adopt whole organ clearing technique that preserves endogenous fluorescent reporters and is compatible with most common imaging platforms, while offering the additional benefit of preserving samples for further downstream imaging analyses.

**Supplementary Figure 1.**
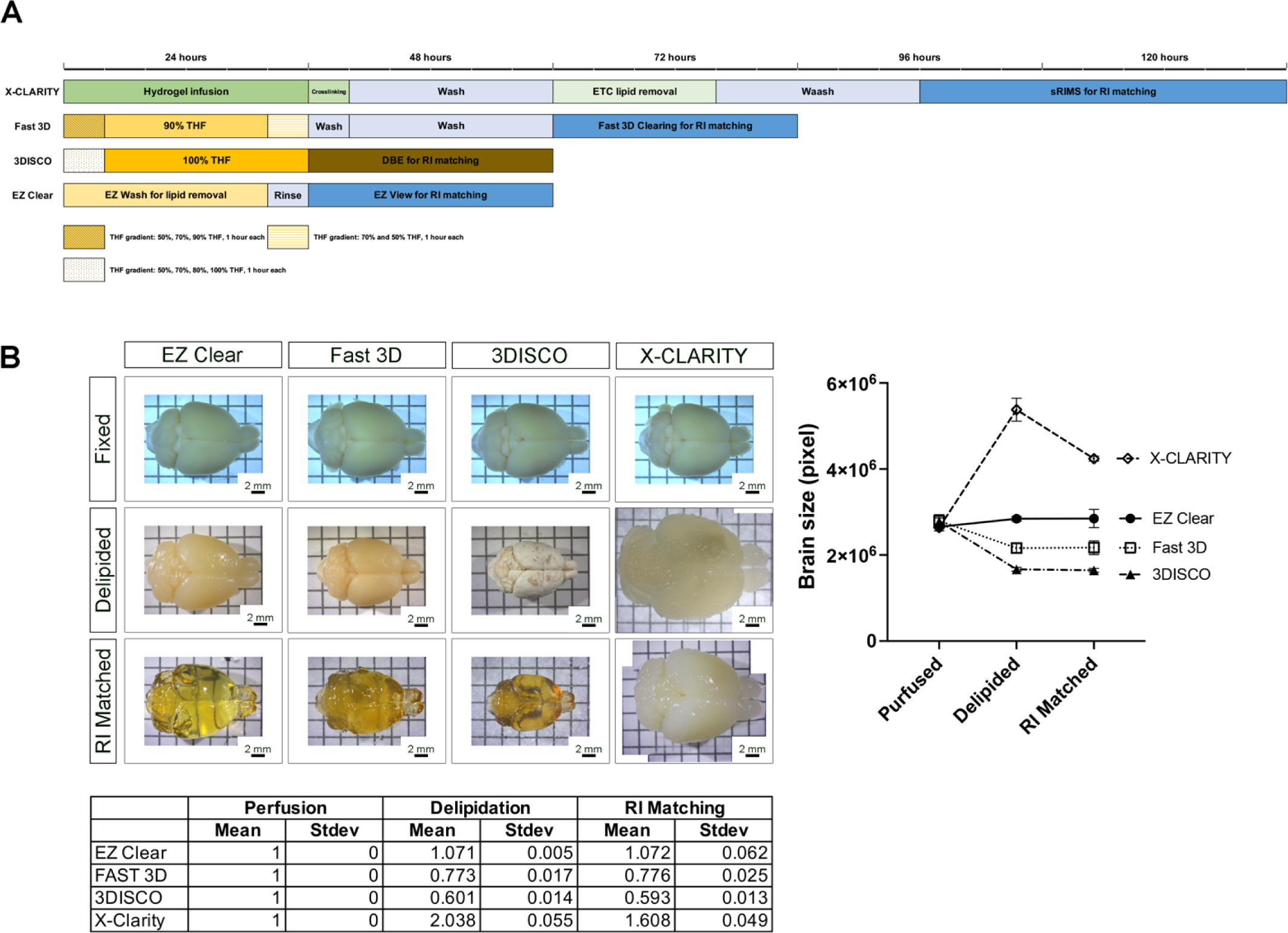
Comparison of EZ Clear to solvent- and aqueous-based clearing methods. (A) Schematic of required steps and time for tissue clearing with X-CLARITY, Fast 3D, 3DISCO, and EZ Clear. (B) Representative images of brains cleared by X-CLARITY, Fast 3D, 3DISCO, and EZ Clear after 4% PFA fixation, delipidation, and RI matching with EZ View (EZ Clear), Fast 3D imaging solution (Fast 3D), DEB (3DISCO), and sRIMS (X-CLARITY). The outlined boundary of the brain from each of the captured brightfield images was used to quantify the total pixel number and thus brain size at each step (n= 4 brains measured for each solution).

**Supplementary Figure 2.**
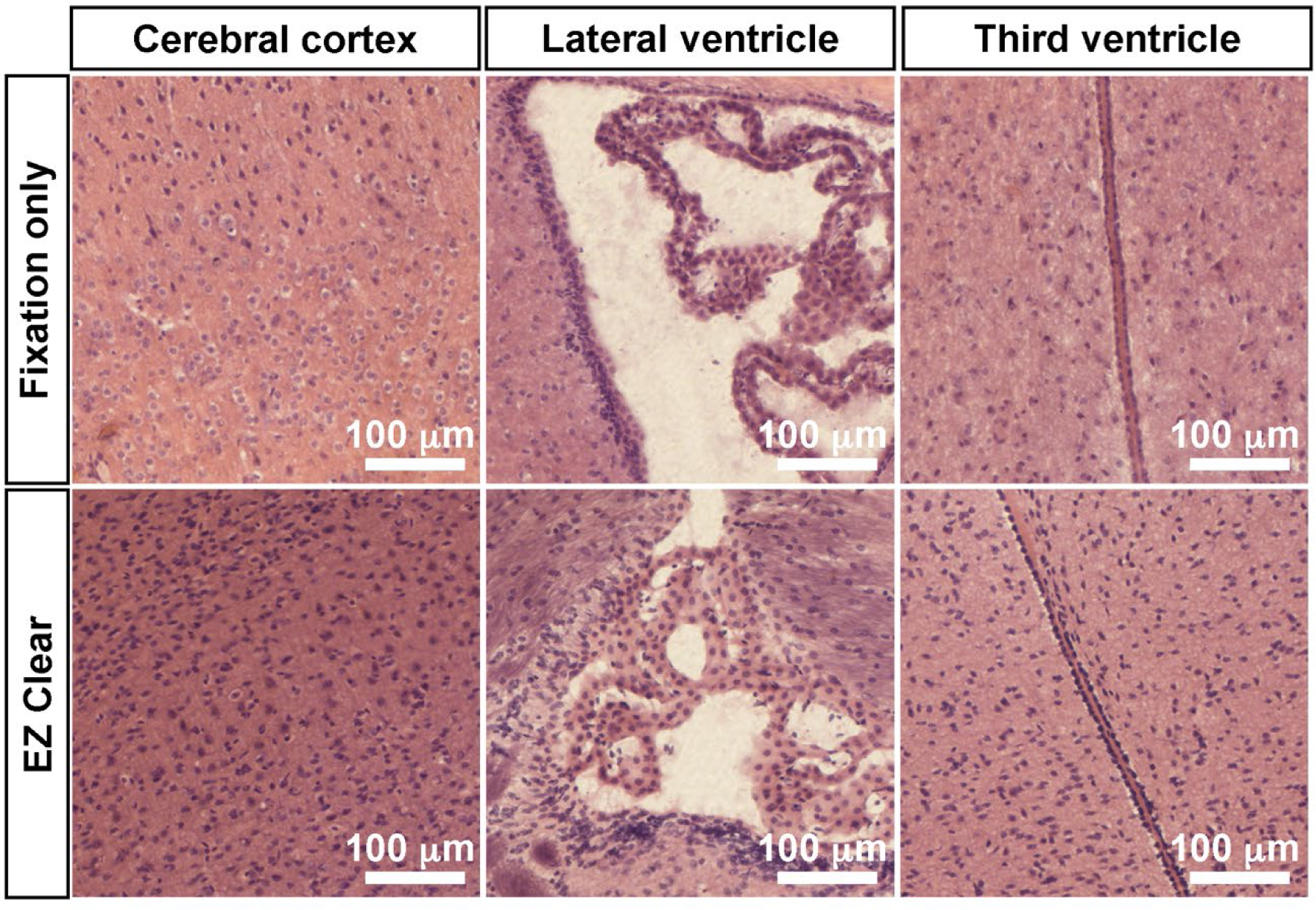
EZ Cleared and imaged samples processed for cryosectioning and H&E staining. (A) Comparison of Hematoxylin and Eosin (H&E) staining of sections from brains that were either fixed or EZ Cleared and lightsheet imaged at cerebral cortex, lateral ventricle, and third ventricle regions.

**Supplementary Figure 3.**
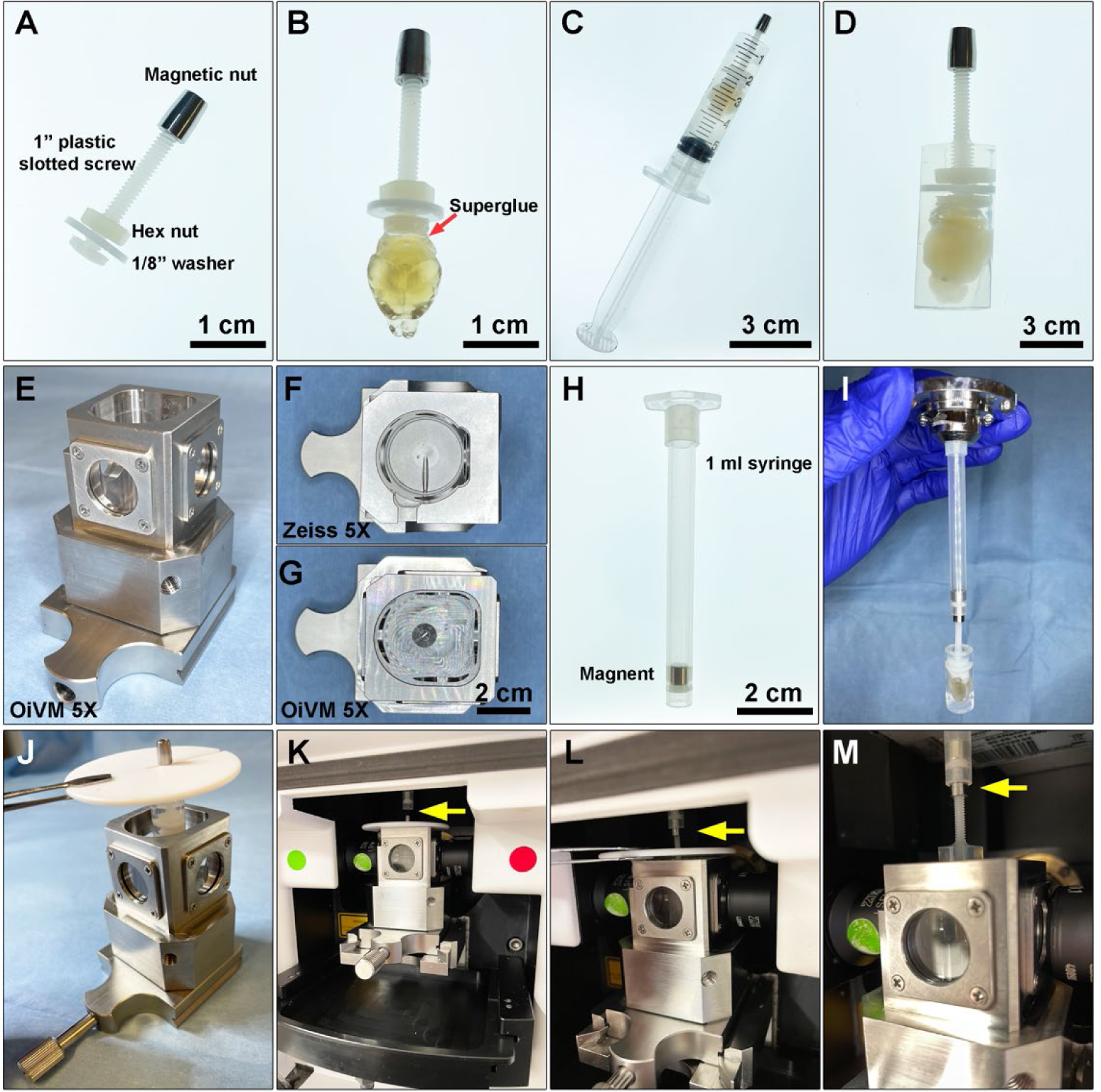
Preparing cleared mouse brain for imaging on Zeiss Lightsheet Z.1. (A) custom designed sample holder with a magnetic nut. (B) Cleared brain sample was attached to the head of the plastic screw with superglue. (C) A 5 ml syringe is used as a casting mold for embedding the cleared brain and the tip of the sample holder in melted 1% agarose. (D) Cleared brain and sample holder embedded in 1% agarose. (E) Custom made OiVM 5X lightsheet chamber. Comparison of the imaging chambers between (F) Zeiss 5X and (G) OiVM 5X. (H) Sample mounting probe with a recess magnet. (I) Demonstration of how the sample mounting probe and sample holder will attach. To load the sample onto lightsheet Z.1, first (J) the embedded sample and holder was placed into the custom-designed imaging chamber by hanging with a PTFE chamber cover with 3 mm opening (Zeiss) by the magnetic nut, then (K) the imaging chamber with sample hanging was assembled onto lightsheet. (L and M) Sample holder was then attached to the probe by lifting the sample with the cover and attached the magnetic nut to the magnet in the probe (yellow arrow). The chamber cover was removed and proceeded with imaging.

**Supplementary Movie 1. Wholemount EZ Cleared *Thy1-EGFP* adult mouse brain imaged by lightsheet fluorescent microscopy**.

**Supplementary Movie 2. Comparison of brain hemispheres perfused with Evans blue and cleared with EZ Clear (right hemisphere) and Fast 3D (left hemisphere)**.

**Supplementary Movie 3. P104 GBM brain perfused with DyLight 649 conjugated *Lycopersicon esculentum* (tomato) lectin (lectin-649)**.

**Supplementary Movie 4. 3D volume rendering of a confocal stack from a GBM brain following cryosectioning and immunostained with GFAP (white) and Hoechst (blue) after EZ Clearing and LFSM imaging of the tumor (green) and blood vessels (red)**.

## ONLINE METHODS

### Materials

All chemicals, reagents, antibodies, instruments, hardware, and software that were used in this study are list in Supplementary Table 1.

### Mice

This study was carried out in strict accordance with the recommendations in the Guide for the Care and Use of Laboratory Animals of the National Institutes of Health. All animal research was conducted according to protocols approved by the Institutional Animal Care and Use Committee (IACUC) of Baylor College of Medicine.

### Generation of endogenous glioma in a mouse model using In Utero electroporation

All mouse CRISPR-IUE GBM gliomas were generated in the CD-1 IGS mouse background. In utero electroporations (IUEs) were performed as previously described ^35^. Briefly, a plasmid containing guide RNAs targeting the tumor suppressor genes *Nf1, Tp53*, and *Pten* was co-electroporated along with a fluorescent reporter EGFP to label tumor cells. Briefly, the uterine horns were surgically exposed in a pregnant dam at E16.5 and the embryos were injected with a DNA cocktail containing the following four plasmids: (1) a single pX330-variant ^37^ construct encoding 3xFlag-NLS-Cas9-NLS, along with three U6 promoter cassettes upstream of validated guide RNA sequences targeting *Nf1* (GCAGATGAGCCGCCACATCGA), *Trp53* (CCTCGAGCTCCCTCTGAGCC) and *Pten* (GAGATCGTTAGCAGAAACAAA) ^38^ (2) a *piggyBac* (PB) helper plasmid with the glial- and astrocyte-specific promoter, *Glast*, driving expression of PB transposase (pGlast-PBase) ^39^, and a PB cargo fluorescent reporter vector (pbCAG-GFP-T2A-GFP or pbCAG-mRFP1). The PBase helper plasmid promotes stable integration of the cargo fluorescent reporter vector, which indelibly labels all descendant cells, allowing one to visualize tumors over time via fluorescence. Following injection of the glioma-inducing CRISPR-Cas9/PB cocktail (2.0 µg/µL pGLAST-PBase, 1.0 µg/µL all other plasmids) into the lateral ventricle of each embryo, embryos were electroporated six times at 100-ms intervals using BTW Tweezertrodes connected to a pulse generator (BTX 8300) set at 33 V and 55 ms per pulse. Voltage was applied across the entire brain to allow uptake of the constructs. The uterine horns were placed back in the cavity, and these dams developed normally, but their electroporated offspring featured malignancies postnatally, as the tumor suppressor deficient cells expanded. CRISPR-IUE tumors were harvested at postnatal day 65 (P65) and P80 from both male and female mice. All mouse experiments were approved by the Baylor College of Medicine Institutional Animal Care and Use Committee.

Animals were perfused at post-natal day 104 (P104) with Lycopersicon esculentum (tomato) lectin fluorescently conjugated with DyLight 649 (Vector Laboratories, Cat. No. DL-1178) before euthanized. Detail procedure for animal perfusion and lectin labeling is listed below. The brains were then dissected out for further processing.

### Animal Perfusion

To perfuse the animal, mouse was deeply anesthetized by CO_2_ inhalation and the chest cavity was opened to expose the beating heart. The right atrium was incised, and the mouse was transcardially perfused through the left ventricle with 10 mL of room temperature 1X PBS, followed by 10 mL of cold 4% PFA. The brain was then dissected from the skull and drop fixed in 4% PFA at 4°C for 24 hours with gentle agitation. After fixation, the brains were washed in 1X PBS 3 times, 30 minutes each at room temperature, then store in 1X PBS with 0.05% sodium azide at 4°C before proceeding with clearing procedure.

For animals that were labeled with lectin, adult mice were injected in their tail vein with 100 μL of Lycopersicon esculentum (tomato) lectin fluorescently conjugated with DyLight 649 (Vector Laboratories, Cat. No. DL-1178). Lectin was allowed to circulate for a minimum of five minutes before tissue was harvested to allow for adequate circulation and binding of the lectin to the endothelium. Mice were then deeply anesthetized by CO_2_ inhalation and the chest cavity was opened to expose the beating heart. An additional 75 μL of fluorescent lectin was injected through the left ventricle using a 31-gauge insulin syringe (BD Biosciences, Cat. No. 324911). Lectin was perfused by hand slowly over one minute and the needle was kept in place for an additional minute after injection to allow the pumping heart to circulate the dye. Afterwards, the right atrium was incised, and the animal was transcardially perfused through the left ventricle with an additional 10 mL of room temperature 1X PBS, followed by 10 mL of cold 4% PFA. The brain was then dissected from the skull and drop fixed in 4% PFA at 4°C for 24 hours with gentle agitation. After fixation, the brains were washed in 1X PBS 3 times, 30 minutes each at room temperature, then store in 1X PBS with 0.05% sodium azide at 4°C before proceeding with clearing procedure.

For animals that were labeled with Evans blue, adult mice were injected in their tail vein with 100 μL of Evans blue solution (2% (w/v) with 0.9% (w/v) NaCl in sterile Milli-Q water, filter sterile with a 0.22 um filter). The dye was allowed to circulate for a minimum of five minutes. Mice were then deeply anesthetized by CO_2_ inhalation and the chest cavity was opened to expose the beating heart. An additional 100 μL of Evans blue solution was injected through the left ventricle using a 31-gauge insulin syringe (BD Biosciences, Cat. No. 324911). The dye was perfused by hand slowly over one minute and the needle was kept in place for an additional minute after injection to prevent dye leaking back out. Allow the injected Evans blue solution to circulate until hindlimbs and tail turns blue, or 5 minutes has reached. The mice were then euthanized with XXX. Dissected out the brain from the skull and drop fixed in 4% PFA at 4°C for 24 hours with gentle agitation. After fixation, the brains were washed in 1X PBS 3 times, 30 minutes each at room temperature, then store in 1X PBS with 0.05% sodium azide at 4°C before proceeding with clearing procedure.

### EZ Clear protocol

Perfused and fixed brain was immersed in 20ml of EZ Wash solution in a glass scintillation vial for lipid removal. EZ Wash solution is consisted of 50% (v/v) tetrahydrofuran (THF, with 250 ppm BHT, Millipore-Sigma 186562) mixed with filtered Milli-Q water. The scintillation vial was covered with foil and placed on an orbital shaker within a vented chemical fume hood, shake gently for 16 hours at room temperature. Removed EZ Wash solution and discarded it carefully in a chemical waste container for proper disposal. Sample was then rinsed in 20ml filtered Milli-Q water 4 times, one hour each to remove any remaining THF. After the last rinse, water was removed as complete as possible, and the vial was left open and vented in the hood for an additional 10 minutes to ensure any remain THF was fully ventilated. At this stage, the delipided samples can be either stored in 1X PBS with 0.05% sodium azide at 4 °C or can be processed immediately for imaging. To render the sample transparent and prepare the sample for imaging, brain was transferred to a 6 ml sample tube. Removed PBS from the vial completely, then 5 ml of EZ View sample mounting and imaging buffer was added into the vial. The EZ View sample mounting and imaging solution is consist of 80% Nycodenz (Accurate Chemical & Scientific 100334-594), 7M of urea, 0.05% sodium azide in 0.02 M phosphate buffer. To prepare the solution, 100 g of Nycodenz, 52.5 g of urea, and 31.25 mg of sodium azide powders were mixed in a 250 ml beaker. 20 ml of 0.02 M of phosphate buffer (pH 7.4) was added and mixed to dissolve the powders using a stir bar overnight. The final volume of the solution was adjusted to 125 ml with additional 0.02 M of phosphate buffer. The dissolved solution was filtered through a vacuum filtration system (Nalgene® vacuum filtration system filter, pore size 0.2 μm, Z370606) and stored at room temperature. The refractive index of the EZ Clear imaging solution was measured on a refractometer (Atago, PAL-RI 3850) and the RI was between 1.512 to 1.518. The sample was protected from light, placed on a shaker, and rocked gently at room temperature for 24 hours to render the sample transparent. All the other mouse organs (eye, heart, lung, liver, kidney, pancreas, testis) were processed with the same procedure.

### 3DISCO clearing

3DISCO clearing was performed according to Erturk et al., 2012 ^5^. Briefly, after perfusion and fixation, adult mouse brain was incubated in graded series of 20 ml of tetrahydrofuran solutions mixed with filtered Milli-Q H_2_O at the concentration of 50, 70, 80, and 100 % (v/v), one hour each, in a scintillation vial at room temperature. The scintillation vial was covered with foil and placed on an orbital shaker within a vented chemical fume hood. Brains were then immersed in fresh 100% THF overnight, followed by 1 hour of another batch of fresh 100% THF. Sample was then immersed in benzyl ether (DBE, Millipore-Sigma, 108014) overnight to render the sample transparent.

### X-CLARITY clearing

X-CLARITY clearing was performed according to manufacturer’s instruction (Logos Biosystems). Briefly, after perfusion and fixation, adult mouse brain was immersed in 5 ml of X-CLARITY Hydrogel Solution with 0.25 % (w/v) of polymerization Initiator VA-044 (Logos Biosystems, C1310X) at 4°C for 24 hours for. The brain infused with the hydrogel solution was then undergo a thermo-induced crosslinking reaction for 3 hours at -90 kPa at 37°C. After the crosslinking, the remaining of the hydrogel solution was removed and brain was washed in 1X PBS 3 times, 1 hour each, then once overnight at 4 °C. Brain was cleared in electrophoretic tissue clearing solution (Logos Biosystems, C13001) using the X-CLARITY Tissue Clearing System set at 0.8 A and 37°C for 15 hours (Logos Biosystems). After electrophoresis, brain was washed in 50 ml of 1X PBS 3 times, 1 hour each, then once overnight at room temperature. To render the tissue transparent for imaging, brains was immersed in sRIMS (70% w/v D-sorbitol in 0.02 M phosphate buffer, pH 7.4) at 4°C until transparent. The refractive index of the sRIMS was measured on refractometer (Atago, PAL-RI 3850) and the RI was between 1.42 to 1.43.

### FAST 3D clearing

FAST 3D clearing was performed according to Kosmidis et al., 2021 ^10^. Briefly, after perfusion and overnight fixation, sample was placed on an orbital shaker at 4 °C, protected from light, and incubated with 20 ml of the following solutions in sequence: (a) 50% (v/v) THF prepared in sterile Milli-Q H_2_O with 20 ul of triethylamine (pH 9.0) (Millipore-Sigma T0886) for 1 hour; (b) 70% THF with 30 ul of triethylamine for 1 hour; and (c) 90% THF with 60 ul of triethylamine overnight. After overnight incubation with 90% THF, samples were then rehydrated with the following solutions: (d) 70% THF with 30 ul of triethylamine for 1 hour followed by (e) 50% THF with 20 ul of triethylamine for 1 hour. Finally, samples were washed with sterile Milli-Q H_2_O 4 times, 10 min each, then washed overnight in sterilized MQ water. To prepare the samples for imaging, the brains were incubated with 4 ml of Fast 3D Clear solution. To prepare the Fast 3D Clear solution, 48 g of Histodenz (Millipore-Sigma, D2158), 0.6 g of Diatrizoic Acid (Millipore-Sigma, D9268), 1.0 g of N-Methyl-D-Glucamine (Millipore-Sigma, M2004), 10 g of Urea, and 0.008 g of sodium azide powders were mixed in a 100 ml beaker. 20 ml of sterile Milli-Q H_2_O was added to dissolve the powder using a stir bar overnight. The final volume of the solution was approximately 50 ml. The dissolved solution was filtered and stored at room temperature. The refractive index of the Fast 3D clear solution was measured on refractometer (Atago, PAL-RI 3850) and the RI was between 1.511 to 1.513. The sample was protected from light, placed on a shaker, and rocked gently at room temperature for 24 hours to render the sample transparent.

### Quantifying changes in brain size

To compare the size changes of brains processed with different clearing protocols, bright field images of the brains were captured with Zeiss Stemi stereo microscope and Labscope Material App at the following stages: after perfusion, lipid removal, and RI matching. For brains that were too large to fit in a single field of view, 4 tiled images with at least 20 % overlap were taken to cover the entire sample and stitched together using Arivis Vision4D. The size change was quantified using Fiji. The “Selection Brush Tool” under “Oval selections” was used to outline the boundary of the brain from the captured brightfield images to measure the total pixel number within the outlined area.

### Sample mounting and whole brain lightsheet imaging

A custom designed sample holder was used for mounting the sample and imaging on a Zeiss Lightsheet Z.1. The sample holder consists of a 1-inch plastic slotted screw (McMaster-Carr, 94690A724) with a nylon hex nut (McMaster-Carr, 94812A200) and a custom-made magnetic nut and a 1/8” washer (Figure S3A). To mount samples, we first applied a small amount of superglue gel (Loctite Gel Control) to the surface of the screw head, and gently pressed the screw head to the brain stem for it to attach (Figure S3B). Then, the luer-lock tip of a 5 ml syringe (BD Bioscience, 309646) was removed using a razor blade, and the remaining syringe was used as a casting mold for mounting the brain. The brain was immersed in 3 mL of 1 % melted agarose prepared in sterile water, and then the solution and immersed brain were gently aspirated to into the sample holder until the agarose solidified completely (Figure S3C). Next, the plunger of the syringe was gently pressed to push the agarose cylinder containing the brain out onto a 10 cm petri dish (Figure S3D). A razor blade was used to trim off extra agarose at the bottom of the cylinder and then the sample and holder were immersed in 25 mL of EZ View imaging solution in a 50 ml conical tube, rocked gently on a horizontal orbital shaker at room temperature (with the conical tube being upright, in a vertical position) overnight to equilibrate.

To image the samples on the Zeiss Lightsheet Z.1, a custom-made, enlarged sample holding chamber was used, which allows sample diameter up to 1 cm compared to original Zeiss 5X chamber (Figure S3E-G). A one mL syringe with a recessed magnet is then used as sample probe to attach to the sample holder (Figure S3H and I). To mount the sample on Lightsheet Z.1 for imaging, the 1 ml syringe with a recessed magnet was first loaded onto the system. Placed the sample into the custom-designed imaging chamber by hanging the mounted sample with a PTFE chamber cover with 3 mm opening (Zeiss) by the magnetic nut (Figure S3J). The imaging chamber with sample hanging was assembled onto lightsheet (Figure S3K). Sample was then attached to the probe by lifting the sample with the chamber cover and attached the magnetic nut to the probe (Figure S3L and M). The PTFE chamber cover was then removed and proceeded with imaging. The samples were imaged with a 5X/0.16 air detection lens and 5X/0.1 illumination lens. The detection lens was zoomed out to 0.5X and the whole brain was acquired at the resolution of 1.829 μm x 1.829 μm x 3.627 μm (X:Y:Z) in tiled sequence with 20 % overlapping. Acquired tiles were aligned and stitched together using Arivis Vision4D.

### Prepare EZ Cleared sample for cryo-sectioning

After the whole brain was imaged in the EZ View solution by LSFM, the sample was equilibrated to PBS by washing in 50 mL of 1X PBS 4 times, 1 hour each, and then once more overnight at room temperature. The brains were then immersed in a 3-step sucrose gradient (10%, 20%, and 30%, prepared in PBS). The sample was incubated at 4 °C for each step until the tissue sank to the bottom of the vial overnight at 4 °C, then embedded in optimal cutting temperature (O.C.T.) medium (Sakura, 4583) and snap frozen on a bed of crushed dry ice, then stored at -80°C until ready for processing. The frozen block was then sectioned on a cryostat (Leica) at 12 μm for H&E staining or at 100 μm for immunostaining.

### Immunostaining

100 mm sections were collected in 1X PBS with 0.05% sodium azide in a 24 well plate (“free floating”). For staining, sections were blocked in 1X PBS + 0.08% Triton X-100 + 2% donkey serum (blocking buffer) overnight at 4 °C, followed by incubation with primary antibodies (CD31 at 1:200, GFAP at 1:200, beta-III tubulin at 1:200) in blocking buffer overnight at 4 °C. Sections were washed 3 times, 1 hour each in 1X PBS, followed by incubation with secondary antibodies (1:500) and Hoechst (Millipore-Sigma,14533, 10 mg/ml stock at 1:1000 dilution) in blocking buffer overnight at 4 °C with gentle agitation on an orbital shaker. Sections stained with SMA-Cy3 (1:200) were stained together with the Hoechst. The next day, after 3, 1-hour washes in 1X PBS at 4 °C, sections were mounted on a charged slide (VWR Micro Slides, 48311-703) with 200 μL of EZ Clear imaging solution and coverslipped with #1 cover glass (VWR Micro Coverglass, 48366-067).

### Histology staining

Slides containing mounted cryosections were equilibrated at room temperature before rehydrating in PBS. They were then subjected to standard Hematoxylin and Eosin staining procedures. Briefly, they were incubated for 2 minutes in Modified Harris Hematoxylin Solution (Millipore-Sigma, HHS32-1L), rinsed with deionized water several times, before incubating for 5 minutes in tap water. They were then dipped 12 times in acid ethanol (0.25% HCl in 70% ethanol), incubated 1 minute in tap water twice and 2 minutes in deionized water. Slides were then incubated in Eosin Y Phloxine B Solution (EMS, 26051-21) for 30 seconds and dehydrated through an ethanol series (70% x 2, 80% x 2, 95% x 2, 99.5% x 2) for 1.5 minutes at each step. Lastly, they were washed in Histoclear II (EMS, 64111-04) twice and coverslipped in DPX new (Millipore-Sigma, HX68428779).

### Statistics analysis

Statistics analysis and one-way ANOVA were performed with GraphPad Prism 9 software.

## ACKNOWLEDGEMENTS

We want to thank Sih-Rong Wu and Dr. Huda Zoghbi for providing us the *Thy1-EGFP* mouse brains and Dr. Nanbing Li-Villarreal for critical review of the manuscript. This project was supported by the Optical Imaging and Vital Microscopy (OiVM) Core for all imaging experiments and the Bioengineering Core at Baylor College of Medicine for manufacturing the custom imaging chamber for the Zeiss Lightsheet Z.1. The authors also want to thank the RNA In Situ Hybridization Core at Baylor College of Medicine, which is, in part, supported by a Shared Instrumentation grant from the NIH (1S10OD016167). This work was supported by grants from the National Institutes of Health (5T32GM088129-10 to W.D.T., R01HL146745 to M.E.D. and J.D.W., R01HD099026 to M.E.D.,

U42OD026645 to M.E.D.), the American Heart Association (19PRE34410104 to M.C.G, 916015 to C.F.S.), the Cancer Prevention Research Institute of Texas (RP200402 to J.D.W.), the Department of Defense (W81XWH-18-1-0350 to J.D.W.) and Canadian Institutes of Health Research (PJT-155922 to J.D.W.).

---

**Supplementary Table 1.** List of resource used in this study.

| Resource | Source | Catalog number |
| --- | --- | --- |
| **Antibodies** |  |  |
| Lycopersicon Esculentum (Tomato) Lectin (LEL, TL), DyLight® 649 | Vector Laboratories | DL-1178-1 |
| Rat Anti-Mouse CD31 | BD Biosciences | 550274 |
| Anti-actin, alpha-smooth muscle – Cy3 | Millipore-Sigma | C6198 |
| GFAP | Invitrogen | 13-0300 |
| Anti-beta III Tubulin | Abcam | ab18207 |
| Donkey Anti-Rat Alexa Fluor 568 | Abcam | Ab175475 |
| Donkey Anti-Rabbit Alexa Fluor 488 | Invitrogen | A21206 |
| Donkey Anti-Rabbit Alexa Fluor 568 | Invitrogen | A10042 |
| **Chemicals and Reagents** |  |  |
| Evans Blue | Millipore-Sigma | E2119 |
| Hoechst | Millipore-Sigma |  |
| Histodenz | Millipore-Sigma | D2158-100g |
| Diatrizoic Acid | Millipore-Sigma | D9268-1g |
| Benzyl ether | Millipore-Sigma | 108014 |
| N-Methyl-D-Glucamine | Millipore-Sigma | M2004-100g |
| Triethylamine | Millipore-Sigma | T0886 |
| Tetrahydrofuran | Millipore-Sigma | 186562 |
| Nycodenz | Accurate Chemical & Scientific | 100334-594 |
| D-Sorbitol | Millipore-Sigma | S1876 |
| Urea | Millipore-Sigma | U5378 |
| Sodium Azide | Millipore-Sigma | S2002 |
| VWR Life Science Agarose I™ | VWR | 0710 |
| Triton X-100 | Fisher scientific | BP-151 |
| Donkey serum | Millipore-Sigma | D9663 |
| Na<sub>2</sub>HPO<sub>4</sub> | Acros Organics | 448140010 |
| NaH<sub>2</sub>PO<sub>4</sub> | Millipore-Sigma | S0751 |
| PBS, 1X solution, pH 7.4, | Fisher scientific | BP24384 |
| Paraformaldehyde | Millipore-Sigma | P6148 |
| Modified Harris Hematoxylin Solution | Millipore-Sigma | HHS32-1L |
| Eosin Y Phloxine B Solution | EMS | 26051-21 |
| Histoclear II | EMS | 64111-04 |
| DPX New | Millipore-Sigma | 100579 |
| Tissue-Tek O.C.T. Compound | Sakura | 4583 |
| **Equipment and Hardware** |  |  |
| Zeiss Lightsheet Z.1 | Zeiss Microscopy | n/a |
| Zeiss LSM 880 with Airyscan FAST Confocal Microscope | Zeiss Microscopy | n/a |
| Zeiss AxioZoom.V16 Stereo/Zoom Microscope | Zeiss Microscopy | n/a |
| Zeiss AxioObserver widefield Microscope | Zeiss Microscopy | n/a |
| Zeiss Stemi Stereo Microscope | Zeiss Microscopy | n/a |
| X-CLARITY™ Tissue clearing system | Logos Biosystems | n/a |
| Pocket Refractometer | Atago | PAL-RI 3850 |
| Plastic Binding Head Slotted Screws, Off-White, 4-40 Thread Size, 1" Long | McMaster-Carr | 94690A724 |
| Micro Slides | VWR | 48311-703 |
| Micro Coverglass | VWR | 48366-067 |
| Nylon Plastic Washer for Number 4 Screw Size, 0.112" ID, 0.206" OD | McMaster-Carr | 90295A340 |
| Nylon Hex Nut, 4-40 Thread Size | McMaster-Carr | 94812A200 |
| Luer-lock, 5 ml syringe | BD Bioscience | 309646 |
| **Software** |  |  |
| Imaris | Bitplane Inc. | Ver. 8.5 |
| Vision4D | Arivis | Ver. 2.12 and 3.5 |
| Zeiss Zen Blue | Zeiss | Ver. 2.6.76.00000 (LSM 880)<br>Ver. 1.1.2.0 (AxioZoom.V16 and AxioObserver.Z1) |
| Zeiss Zen Black | Zeiss | Ver. 9.2.5.54 (Lightsheet Z.1)<br>Ver. 14.0.23.201 (LSM 880) |
| ImageJ/Fiji | NIH | Ver. 1.53c |
| Prism | GraphPad | Ver. 9.2.0 |

